# Opioid antagonism reduces wanting by strengthening frontostriatal connectivity

**DOI:** 10.1101/2021.06.20.449203

**Authors:** Alexander Soutschek, Susanna C. Weber, Thorsten Kahnt, Boris B. Quednow, Philippe N. Tobler

**Affiliations:** Department of Psychology, Ludwig Maximilian University, Munich, Germany; Zurich Center for Neuroeconomics, Department of Economics, University of Zurich, Zurich, Switzerland; Department of Neurology, Northwestern University Feinberg School of Medicine, Chicago, Illinois, United States of America; Experimental and Clinical Pharmacopsychology, Department of Psychiatry, Psychotherapy and Psychosomatics, Psychiatric Hospital, University of Zurich, Zurich, Switzerland; Neuroscience Center Zurich, University of Zurich and Swiss Federal Institute of Technology Zurich, Zurich, Switzerland

**Author notes:** Correspondence should be addressed to: Alexander Soutschek, Ludwig Maximilian University Munich, Department of Psychology, Leopoldstr. 13, 80802 Munich, Germany. Author contributions: SCW, TK, BBQ, and PNT designed research; SCW conducted research; AS and SCW analyzed data; AS and PNT wrote manuscript; SCW, TK, and BBQ approved final manuscript version.

**Keywords:** reward, wanting, liking, dopamine, opioid, amisulpride, naltrexone, dorsolateral prefrontal cortex, striatum

## Abstract

Goal-directed behavior depends on both motivational (“wanting”) and hedonic (“liking”) dimensions of rewards. Previous animal and human research linked wanting and liking to anatomically and neurochemically distinct brain mechanisms, but it remains unknown as to how the different brain regions and neurotransmitter systems interact in processing distinct reward dimensions. Here, we assessed how pharmacological manipulations of opioid and dopamine receptor activation modulate the neural processing of wanting and liking in humans in a randomized, placebo-controlled, double-blind clinical trial. Blocking opioid receptor activation with naltrexone selectively reduced wanting of rewards, which on a neural level was reflected by stronger coupling between dorsolateral prefrontal cortex and the striatum under naltrexone compared with placebo. Our findings thus provide insights into how opioid receptors mediate frontostriatal gating of specifically motivational, not hedonic, aspects of rewards.

## Introduction

Rewards are central for goal-directed behavior as they induce approach behavior towards valued outcomes (Schultz, 2015). Theoretical models distinguish between behavioral dimensions of rewards, such as the motivational drive to obtain rewards (“wanting”) versus the hedonic pleasure associated with reward consumption (“liking”) (Berridge, 1996; Berridge & Kringelbach, 2015; Berridge, Robinson, & Aldridge, 2009). Dysfunctions in wanting and liking of rewards belong to the core symptoms to addiction, which can be conceptualized as a wanting-dominated state with deficits in switching to liked non-drug rewards (Berridge et al., 2009). It is thus important to obtain a better understanding of the human brain mechanisms underlying wanting and liking. Previous animal research suggested that wanting and liking relate to dissociable neurochemical mechanisms: Dopaminergic activity is thought to modulate the wanting component of rewards, but not liking (Berridge & Valenstein, 1991). In contrast, the opioidergic system has later been associated with both wanting and liking (Berridge & Kringelbach, 2015). Human studies support the hypothesized link between dopaminergic activation and cue-triggered wanting (Hebart & Gläscher, 2015; Soutschek, Kozak, et al., 2020; Weber et al., 2016) as well as the motivation to work for rewards (Chong et al., 2015; Korb et al., 2020; Skvortsova, Degos, Welter, Vidailhet, & Pessiglione, 2017; Soutschek, Gvozdanovic, et al., 2020; Westbrook et al., 2020; Zénon, Devesse, & Olivier, 2016). Noteworthy, one of these studies suggests that dopamine changes only implicit measures of wanting (motivation to work for rewards), but not self-report wanting ratings (Korb et al., 2020).

Consistent with animal findings, pharmacological manipulations of the opioid system affected both wanting and liking aspects of rewards in humans (C. Buchel, S. Miedl, & C. Sprenger, 2018; Christian Buchel, Stephan Miedl, & Christian Sprenger, 2018; Chelnokova et al., 2014; Eikemo et al., 2016). Less is known, however, about the neuroanatomical basis of human wanting and liking. Both dopaminergic and opioidergic neurons project to reward circuits in the striatum as well as to the prefrontal cortex, and recent neuroimaging findings suggest that these regions indeed play a role in processing of wanting and liking (Weber, Kahnt, Quednow, & Tobler, 2018). In particular, the ventral striatum encodes the currently behaviorally relevant reward dimension and dynamically switches functional connectivity with wanting- and liking-encoding prefrontal regions accordingly (frontostriatal gating hypothesis). Thus, processing of wanting and liking appears to be dissociable on both a neurochemical and an anatomical basis. However, it remains unknown how pharmacological and connectivity-related brain mechanisms interact. We investigated whether frontostriatal connectivity gating of motivational and hedonic judgments is orchestrated by dissociable neurotransmitter systems.

To test this hypothesis, the current study investigated the impact of pharmacologically manipulating dopaminergic and opioidergic systems on the neural processing of wanting and liking information. Specifically, we administered a task that allows distinguishing between wanting and liking dimensions of valued goods and assessed how pharmacologically blocking dopaminergic (using the dopamine antagonist amisulpride) or opioidergic neurotransmission (with the opioid antagonist naltrexone) changes the frontostriatal gating of parametric wanting and liking judgments. We hypothesized that, on a behavioral level, opioid receptor blockade will reduce both wanting and liking, whereas reduced dopaminergic neurotransmission will selectively affect wanting. We further hypothesized that on a neural level the behavioral effects of the pharmacological manipulations are mirrored by changes in the frontostriatal gating of wanting and liking information. In particular, we expected that the frontostriatal gating of wanting will be orchestrated by opioidergic and dopaminergic activation (as these neurotransmitters have been related to the processing of wanting), whereas frontostriatal gating of liking should be reduced after blockade of opioidergic neurotransmission.

## Results

### Opioid antagonism reduces wanting ratings

We analyzed the data of healthy young volunteers who rated how much they wanted or liked everyday items in the MRI scanner. We collected wanting and liking ratings for all items twice, once before (pre-test session) and once after (post-test session) participants played a game on the computer where they randomly won or lost 50% of the items. This allowed us to assess whether participants behaviorally distinguished between wanting and liking ratings, because based on our previous findings we expected that winning and losing items has dissociable effects on wanting and liking (Weber et al., 2018). To test the impact of pharmacologically manipulating dopaminergic and opioidergic receptor activation on wanting and liking, participants received either naltrexone (N=37), amisulpride (N=40), or placebo (N=39) prior to performing the task in the scanner.

First, we performed a sanity check whether participants distinguished between wanting and liking ratings by assessing the impact of winning versus losing items on wanting and liking ratings in the post-test session. As recommended for pre-test/post-test designs (Dugard & Todman, 1995), we regressed ratings in the post-test session on item-specific pre-test ratings. Moreover, we included predictors for *Judgement* (wanting versus liking), *Item type* (lost versus won), and the interaction terms. As expected based on our previous findings (Weber et al., 2018), we observed a marginally significant *Judgement* × *Item type* interaction, β = 0.50, *t*(111) = 1.61, *p* = 0.05, one-tailed, suggesting that winning versus losing items had dissociable effects on wanting versus liking of the items (Figure 1B and Table 1). Separate analyses for wanting and liking ratings revealed that wanting ratings did not differ between won and lost items, β = 0.29, *t*(115) = 0.64, *p* = 0.52, whereas liking was more strongly reduced for lost than for won items, β = 0.92, *t*(1016) = 3.29, *p* = 0.001. Participants thus behaviorally distinguished between wanting and liking ratings.

**Table 1.** Results of MGLM-1 on wanting and liking ratings in the post-test as function of *Judgement* (wanting versus liking), *Item type* (lost versus won), and *Pre-test* ratings. Standard errors of the mean (SE) are in brackets.

|  | **beta (SE)** | **t-value** | **df** | **p-value** |
| --- | --- | --- | --- | --- |
| Intercept | 0.21 (0.66) | 0.32 | 119 | 0.75 |
| Judgement | -2.27 (0.64) | 3.54 | 125 | <0.001 |
| Item type | 0.07 (1.10) | 0.07 | 98 | 0.95 |
| Pre-test | 79.87 (0.50) | 158.30 | 124 | <0.001 |
| Judgement × Item type | 2.01 (1.24) | 1.62 | 109 | 0.10* |
| Judgement × Pre-test | 2.41 (0.61) | 3.93 | 289 | <0.001 |
| Item type × Pre-test | -1.62 (0.71) | 2.26 | 133 | 0.03 |
| Judgement × Item type × Pre-test | 0.91 (0.90) | 1.01 | 288 | 0.31 |

\* $p = 0.05$ with one-tailed testing, directed apriori hypothesis based on (Weber et al., 2018)

**Figure 1.**
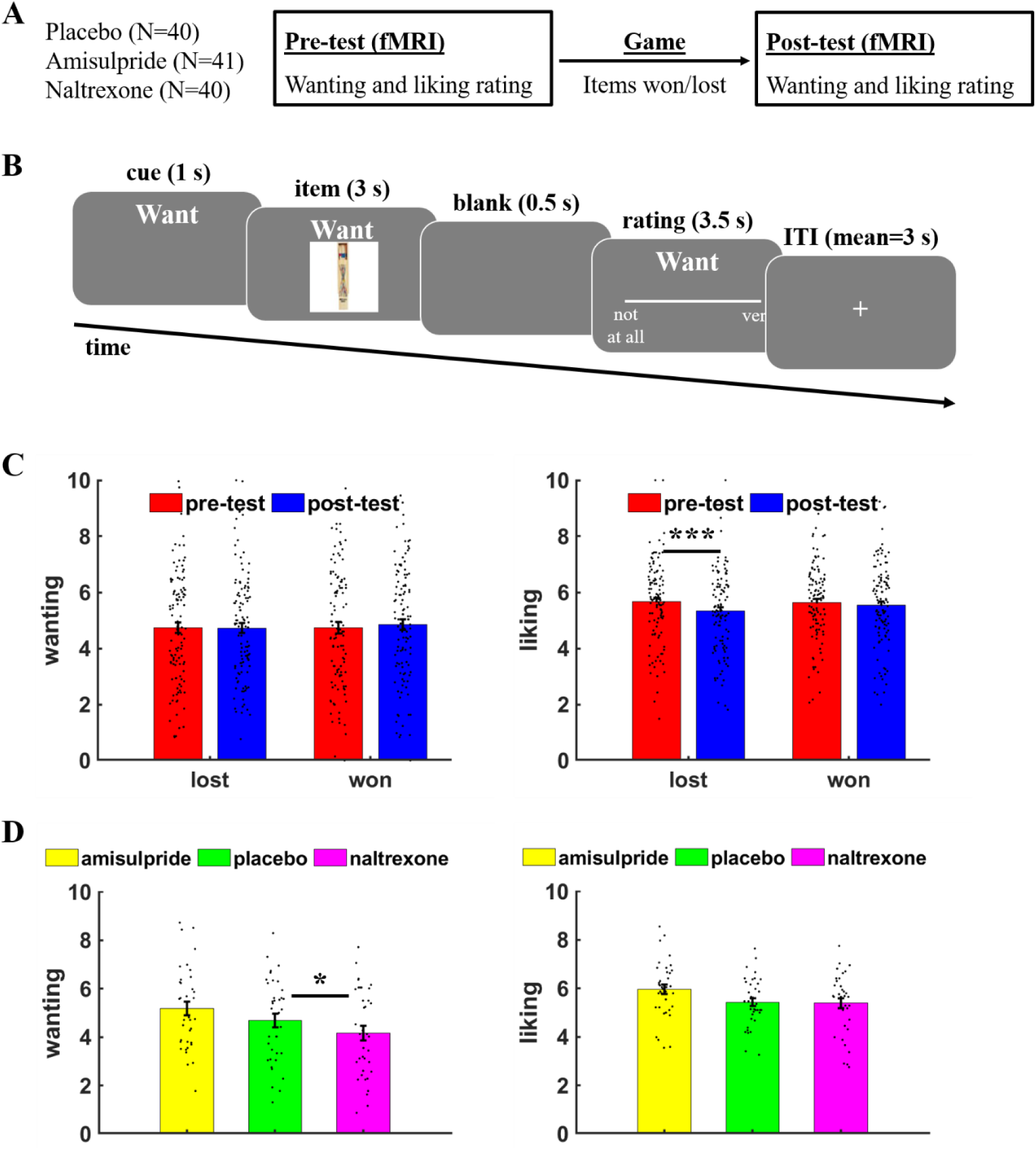
Task procedure and behavioral results. (A) Participants rated how much they wanted or liked objects before (pre-test) or after (post-test) they could win or lose these items in a game between the scanning sessions. (B) On each trial, a cue indicated whether a wanting or liking rating was required, followed by the presentation of the current object (here: a pick-up sticks game). Participants had to rate how much they wanted or liked the presented object within 3.5s, then the next trial started after a variable inter-trial interval (mean = 3s). (C) Liking ratings were significantly reduced for objects that were lost relative to won in the gamble outside the scanner, while wanting ratings did not significantly differ between lost versus won items. (D) The opioid antagonist naltrexone significantly reduced wanting ratings relative to placebo, while liking ratings were unaffected by naltrexone or the dopamine antagonist amisulpride. For illustration purposes, wanting/liking ratings are plotted on a scale from 0 to 10. Error bars indicate standard error of the mean, black dots represent individual data points. *\** p < 0.05, *** *p* < 0.001.

Next, we assessed the impact of dopamine and opioid receptor blockade on wanting and liking judgements. We analyzed ratings (pre- and post-test) with predictors for *Amisulpride* (versus placebo), *Naltrexone* (versus placebo), *Judgement, Session* (pre-test versus post-test), and the interaction terms. This analysis provided evidence that blocking opioid neurotransmission differentially affected wanting and liking ratings, *Naltrexone* × *Judgement*, β = 7.02, *t*(125) = 2.36, *p* = 0.02, while we observed no significant effects for amisulpride, β = 3.79, *t*(126) = 1.30, *p* = 0.20 (Figure 1C and Table 2). Judgement type-specific analyses suggested that wanting ratings were significantly reduced under naltrexone relative to placebo, β = −13.85, *t*(115) = 2.12, *p* = 0.04, whereas amisulpride did not change wanting ratings relative to placebo, β = −1.39, *t*(116) = 0.22, *p* = 0.83. Neither naltrexone nor amisulpride showed significant effects on liking, both *t* < 1.17, both *p* > 0.24. Taken together, our findings provide evidence for involvement of opioidergic neurotransmission in wanting judgements.

**Table 2.** Results for MGLM-2 assessing drug effects on wanting and liking ratings as function of *Drug* (amisulpride versus placebo and naltrexone versus placebo), *Judgement* (wanting versus liking), and *Session* (pre-test versus post-test). Standard errors of the mean (SE) are in brackets.

|  | **beta (SE)** | **t-value** | **df** | **p-value** |
| --- | --- | --- | --- | --- |
| Intercept | 3.62 (5.50) | 0.66 | 98 | 0.51 |
| Amisulpride | 2.37 (5.09) | 0.47 | 114 | 0.64 |
| Naltrexone | -6.80 (5.19) | 1.31 | 114 | 0.19 |
| Judgement | 4.38 (2.08) | 2.11 | 125 | 0.04 |
| Session | -3.30 (1.98) | 1.67 | 1612 | 0.10 |
| Amisulpride × Judgement | 3.79 (2.92) | 1.30 | 126 | 0.20 |
| Naltrexone × Judgement | 7.02 (2.98) | 2.36 | 125 | 0.02 |
| Amisulpride × Session | 0.02 (2.79) | 0.00 | 1629 | 0.99 |
| Naltrexone × Session | 2.70 (2.84) | 0.95 | 1618 | 0.34 |
| Judgement × Session | -1.09 (1.97) | 0.55 | 1845 | 0.58 |
| Amisulpride × Judgement × Session | -1.50 (2.78) | 0.54 | 1856 | 0.59 |
| Naltrexone × Judgement × Session | -2.95 (2.83) | 1.04 | 1853 | 0.30 |

### Opioid antagonism reduces frontostriatal gating of wanting

Next, we investigated the neural mechanisms underlying the impact of opioid antagonism on wanting. Following the procedures from our previous study (Weber et al., 2018), we first determined the neural correlates of wanting and liking by computing GLM-1 in which onset regressors for wanting and liking judgements were modulated by non-orthogonalized parametric modulators for wanting and liking ratings. Wanting ratings (independently of the required judgement type) correlated with activation in ventromedial prefrontal cortex (VMPFC; *z* = 7.32, whole-brain FWE-corrected, *p* < 0.001, peak = [0 44-7]), dorsolateral prefrontal cortex (DLPFC; *z* = 6.63, whole-brain FWE-corrected, *p* < 0.001, peak = [−21 38 44]), and posterior cingulate cortex (PCC; *z* = 5.29, whole-brain FWE-corrected, *p* = 0.002, peak = [−3 37 38]) (Figure 2A and Table 3). Liking ratings correlated with BOLD signal changes in dorsal PCC (*z* = 4.37, whole-brain FWE-corrected, *p* = 0.02, peak = [−9 −64 38]) (Figure 2B and Table 4). Moreover, we also replicated our previous finding that liking ratings correlate with activity in orbitofrontal cortex (OFC) when applying small-volume correction (SVC; anatomical mask for the OFC based on the wfupickatlas; *z* = 3.20, small volume FWE-corrected, *p* = 0.046, peak = [−21 50 −4]). Together, these data replicate our previous findings that wanting and liking are correlated with activation in VMPFC and OFC, respectively. However, we observed no significant effects of naltrexone or amisulpride (relative to placebo) on these neural representations of wanting or liking in these regions (or at whole-brain level), even at lenient statistical thresholds (*p* < 0.001 uncorrected, cluster size > 20 voxels).

**Table 3.**
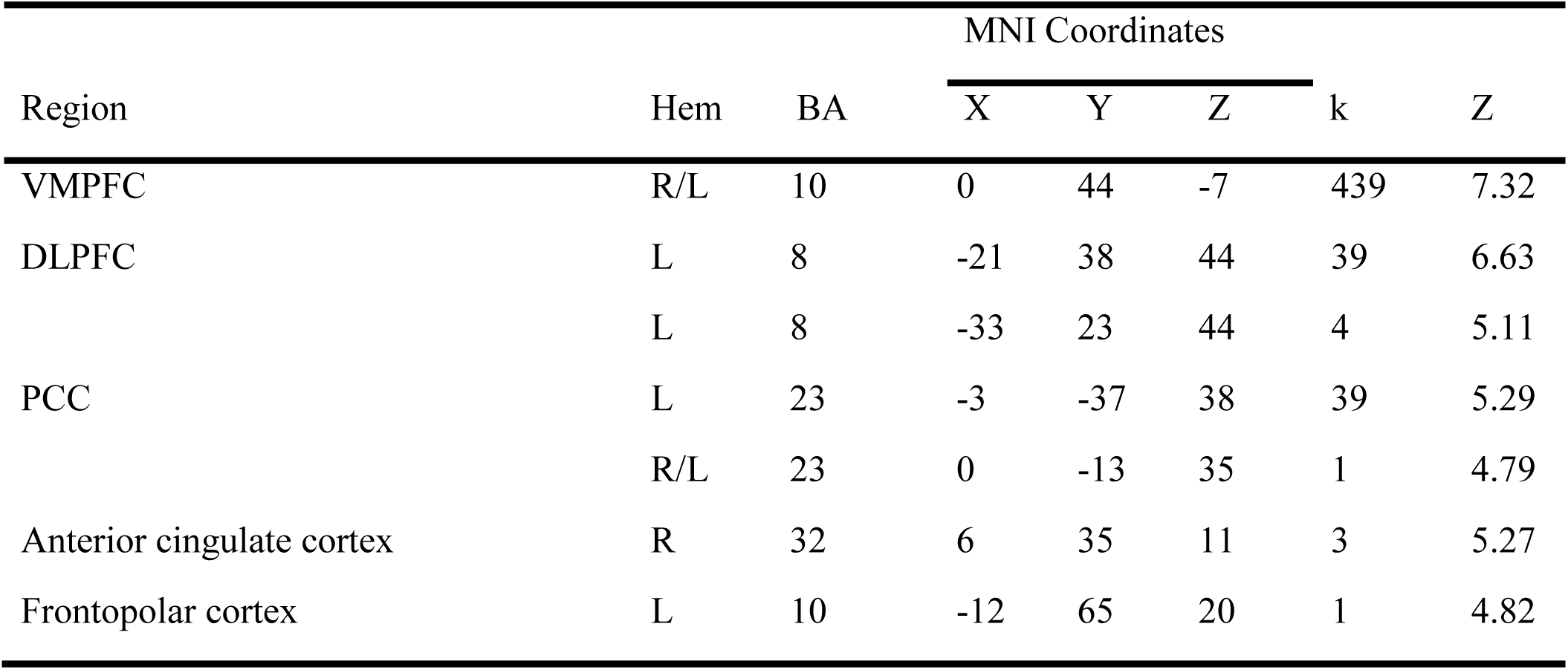
Anatomical locations and MNI coordinates of the peak activations correlating with wanting ratings in GLM-1. We report activations surviving whole-brain FWE correction at peak level (*p* < 0.05). Hem = Hemisphere (L = left, R = right); BA = Brodmann area

| Region | Hem | BA | MNI Coordinates |  |  | k | Z |
| --- | --- | --- | --- | --- | --- | --- | --- |
|  |  |  | X | Y | Z |  |  |
| VMPFC | R/L | 10 | 0 | 44 | -7 | 439 | 7.32 |
| DLPFC | L | 8 | -21 | 38 | 44 | 39 | 6.63 |
|  | L | 8 | -33 | 23 | 44 | 4 | 5.11 |
| PCC | L | 23 | -3 | -37 | 38 | 39 | 5.29 |
|  | R/L | 23 | 0 | -13 | 35 | 1 | 4.79 |
| Anterior cingulate cortex | R | 32 | 6 | 35 | 11 | 3 | 5.27 |
| Frontopolar cortex | L | 10 | -12 | 65 | 20 | 1 | 4.82 |

**Table 4.**
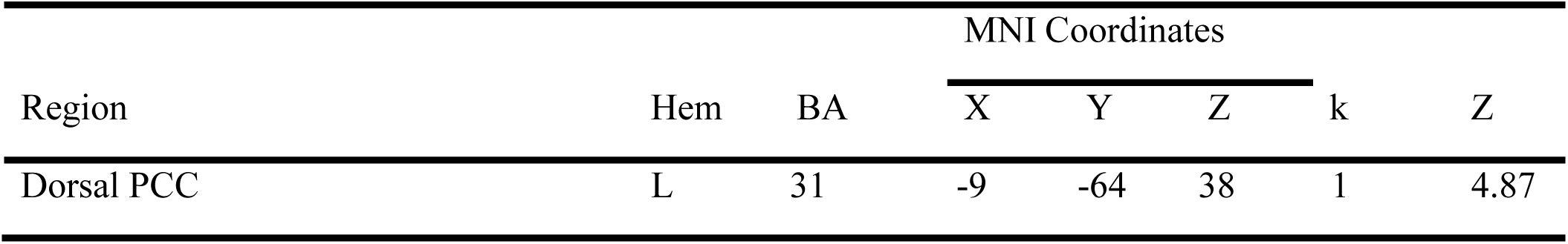
Anatomical locations and MNI coordinates of the peak activations correlating with liking ratings in GLM-1. We report activations surviving whole-brain FWE correction at peak level (*p* < 0.05). Hem = Hemisphere (L = left, R = right); BA = Brodmann area

**Figure 2.**
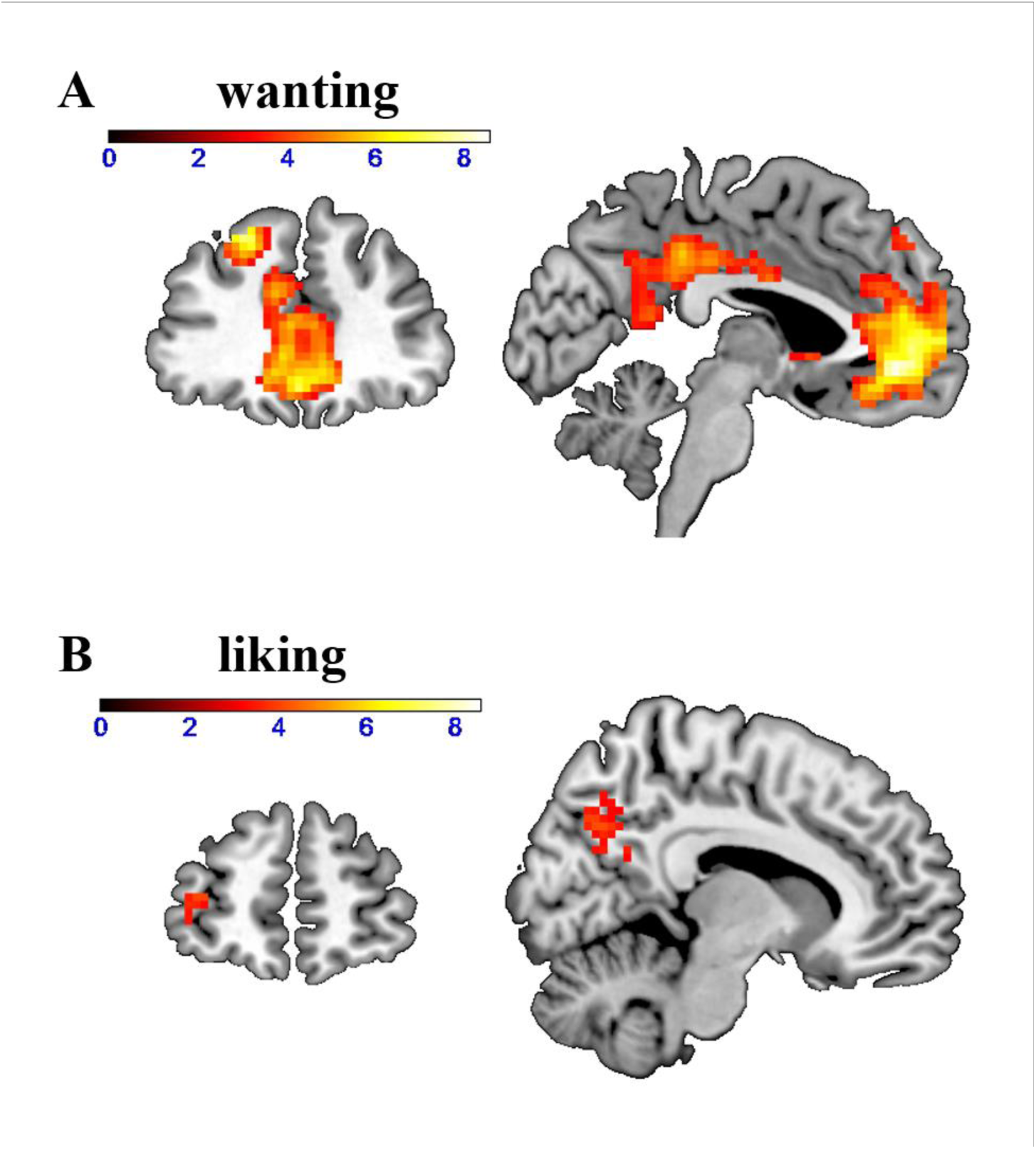
Neural correlates of (A) wanting and (B) liking independently of behavioral relevance. Wanting correlated with activation in DLPFC, VMPFC, and PCC (whole-brain FWE-corrected). Liking correlated with activation in dorsal PCC (whole-brain FWE-corrected) and orbitofrontal cortex (small-volume FWE-corrected).

Previous research showed that wanting-related prefrontal activation is functionally coupled with the ventral striatum depending on the behavioral relevance of wanting judgements (Weber et al., 2018). Consistent with our previous finding, striatal activation was significantly correlated with wanting ratings when those where behaviorally relevant (wanting ratings on wanting trials in GLM-2), *z* = 4.46, *p* = 0.003, peak = [−6 11 −1], small volume FWE-corrected with anatomical mask for the striatum. We therefore assessed whether wanting-related prefrontal regions are functionally connected with the striatum by conducting a PPI analysis with the striatum as seed region. To test whether the pharmacological manipulations changed the functional connectivity of the striatum with specifically wanting-related brain regions, we applied small-volume correction using mask including the significant wanting-correlated voxels in DLPFC and VMPFC in GLM-1 (thresholded with FWE at peak level, k = 478). On wanting trials, we observed enhanced functional coupling between striatum and DLPFC as a function of increasing wanting ratings (wanting ratings on wanting trials: *z* = 3.87, small volume FWE-corrected, *p* = 0.02, peak = [−21 41 41]). Moreover, DLPFC-striatum connectivity on wanting trials was stronger for wanting than for liking ratings (wanting > liking ratings on wanting trials: *z* = 3.64, small volume FWE-corrected, *p* = 0.04, peak = [−21 41 41]) (Figure 3A). The wanting-dependent DLPFC-striatum coupling is consistent with previous findings that connectivity between the stratum and prefrontal correlates of wanting depends on whether wanting judgements are behaviorally relevant (Weber et al., 2018).

**Figure 3.**
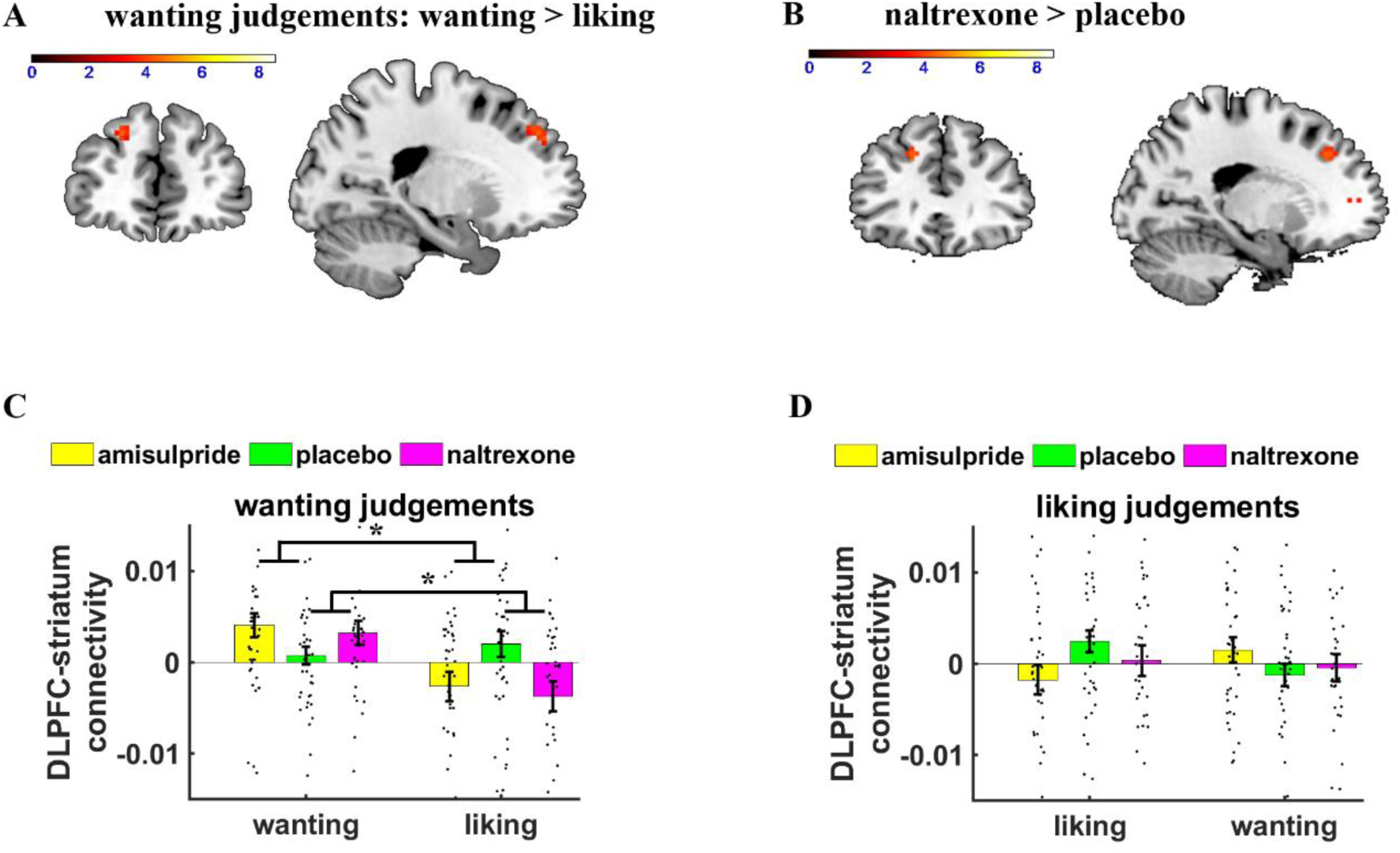
Effects of Judgment type and drug on parametric striatal connectivity. (A) On wanting trials (collapsed across drug groups), DLPFC-striatum connectivity was enhanced for wanting relative to liking aspects of rewards. (B) Wanting-related DLPFC-striatum coupling was significantly stronger under naltrexone compared with placebo. (C, D) Extracted parameter estimates for DLPFC (as defined by the significant cluster in GLM-1), separately for wanting and liking judgements. (C) If wanting judgements were behaviorally relevant, naltrexone increased wanting-relative to liking-related DLPFC-striatum connectivity. (D) No significant drug effects on DLPFC-striatum connectivity were observed on liking trials. Error bars indicate standard error of the mean, black dots represent individual data points. *\** p < 0.05.

Next, we tested how our pharmacological manipulation changed functional connectivity between the striatum and wanting-related cortical regions. Compared with placebo, naltrexone increased DLPFC-striatum coupling for wanting relative to liking ratings on wanting trials (naltrexone > placebo for wanting > liking ratings on wanting trials: *z* = 3.81, small volume FWE-corrected, *p* = 0.02, peak = [−18 35 38]) (Figure 3B). Moreover, the impact of naltrexone on DLPFC-striatum connectivity was significantly stronger on wanting than on liking trials, (“(wanting > liking ratings)_wanting trials_ > (wanting > liking ratings)_liking trials_” -related connectivity in the naltrexone relative to the placebo group: *z* = 3.55, small volume FWE-corrected, *p* = 0.05, peak = [−18 35 38]). Thus, the effects of naltrexone on frontostriatal connectivity were specific for wanting judgements. We observed no further regions showing significantly reduced wanting-related connectivity under naltrexone relative to placebo, and we also observed no significant differences between amisulpride and placebo as well as naltrexone and amisulpride. Thus, blocking opioid neurotransmission strengthened the functional connection between the striatum and prefrontal cortex when wanting judgements were behaviorally relevant.

For completeness, despite having observed no significant drug effects on liking in the behavioral analysis, we also performed whole-brain analyses assessing which brain regions show enhanced functional connectivity as function of liking ratings. On liking trials, connectivity with the striatum was stronger for liking than for wanting ratings in OFC, (*z* = 3.16, small volume FWE-corrected, *p* = 0.05, peak = [−33 50 −7]), replicating our previous findings. However, no brain regions showed significant effects of naltrexone or amisulpride (relative to placebo) on liking-related connectivity with the striatum even at low, exploratory statistical thresholds (*p* < 0.001, cluster size > 20 voxels).

To assess the robustness of the naltrexone effects on wanting-related DLPFC-striatum connectivity, we extracted parameter estimates from the significant wanting-related DLPFC cluster in GLM-1 and regressed the parameter estimates on predictors for Naltrexone, Amisulpride, Judgement, Relevance, and the interaction terms (using the lmer function in R). The significant *Naltrexone* × *Judgement* × *Relevance* interaction, β = 6.0e-03, *t*(452) = 2.46, *p* = 0.01, replicated the finding that naltrexone had dissociable effects on wanting and liking as a function of the behavioral relevance of these reward components. We also observed a significant *Amisulpride* × *Relevance* interaction, β = 4.1e-03, *t*(452) = 2.42, *p* = 0.02. Separate analyses for wanting and liking judgements revealed that in wanting trials DLPFC-striatum connectivity was enhanced for wanting compared with liking ratings under both naltrexone, *Naltrexone* × *Relevance* interaction, β = 5.6e-03, *t*(226) = 3.19, *p* = 0.002, and amisulpride, *Amisulpride* × *Relevance* interaction, β = 4.1e-03, *t*(226) = 2.35, *p* = 0.02 (Figure 3C). In contrast, liking judgements showed no significant effects of naltrexone or amisulpride relative to placebo, all *t* < 1.34, all *p* > 0.18 (Figure 3D). This result supports our findings based on SVC according to which naltrexone increases wanting-related relative to liking-related DLPFC-striatum connectivity on wanting trials, and hints at a similar function for amisulpride (though this was not evident in the SVC-based analysis).

## Discussion

In animal models, the wanting and liking dimensions of rewards have been shown to be processed by distinct brain regions and neurotransmitter systems, but in humans it remained unclear so far how opioidergic and dopaminergic systems orchestrate the processing of wanting and liking. The current findings provide evidence for dissociable roles of opioidergic neurotransmission in processing the two dimensions of rewards on both a behavioral and a neural level. Behaviorally, blocking opioidergic activation with naltrexone selectively reduced wanting, not liking, ratings for non-consumable goods. On a neural level, this reduction in wanting was reflected by changes in DLPFC-striatum connectivity: When wanting judgements were required, DLPFC-striatum connectivity was significantly stronger for the behaviorally relevant wanting ratings than for the irrelevant liking dimension of rewards.

Importantly, wanting-related functional coupling between DLPFC and striatum was significantly stronger under naltrexone than under placebo. This is consistent with recent findings relating opioid receptor blockade with increased connectivity between the prefrontal control system and reward circuits (Elton, Dove, Spencer, Robinson, & Boettiger, 2019; Lim et al., 2019) and suggesting prefrontal kappa opioid receptors to mediate the impact of naltrexone on drug craving in alcohol use disorder (de Laat et al., 2019). Through corticostriatal loops, the striatum receives input from several cortical regions and can prioritize processing of behaviorally relevant information (Frank, 2011). DLPFC provides inhibitory input to the striatum and was shown to reduce wanting-related activation in the striatum (Dong et al., 2020; Koob & Volkow, 2010), consistent with the importance of frontostriatal loops for self-control (van den Bos, Rodriguez, Schweitzer, & McClure, 2014). Opioid receptor blockade might thus reduce wanting of rewards by strengthening inhibitory input from DLPFC to the neural reward system.

Contrary to our hypotheses, we did not observe effects of naltrexone on liking or amisulpride effects on wanting. Interestingly, however, a recent study observed no influences of opioid and dopamine antagonists on explicit wanting and liking ratings but only on implicit measures of these reward dimensions (Korb et al., 2020). In fact, previous studies reporting effects of dopaminergic manipulations on wanting operationalized wanting via implicit measures rather than explicit ratings (Soutschek, Gvozdanovic, et al., 2020; Soutschek, Kozak, et al., 2020; Weber et al., 2016), while studies using explicit ratings observed no or only weak effects of pharmacological manipulations (Case et al., 2016; Ellingsen et al., 2014; Løseth, Eikemo, & Leknes, 2019). Moreover, given that in our study participants had to provide liking ratings without being able to actually consume or handle the items, our measurements might have been less sensitive than those of other studies assessing the liking of consumed rewards (C. Buchel et al., 2018; Chelnokova et al., 2014; Eikemo et al., 2016). It is also worth noting that it has recently been suggested that amisulpride shows, if any, only weak effects on BOLD signal changes in the reward system (Grimm et al., 2020). One should thus be careful with interpreting these unexpected null findings as being inconsistent with previous pharmacological results manipulating dopaminergic activity with different compounds than amisulpride.

Interestingly, the ROI analysis provided some evidence for amisulpride effects on wanting at the neural level, as amisulpride increased wanting-related DLPFC-striatum connectivity, similar to the findings for naltrexone. However, the impact of amisulpride on frontrostriatal gating of wanting needs to be interpreted with caution, given the lack of significant amisulpride effects on behavior.

Our findings have important implications for clinical research, given that dysfunctions in wanting and liking are prevalent in several psychiatric disorders. Substance use disorders, for example, are characterized by increased wanting of drugs as reflected in craving symptoms (Berridge, 2012; Edwards, 2016), and craving has been linked to impairments in prefrontal top-down control over the striatum (Feil et al., 2010). Naltrexone is approved in several countries for the treatment of alcohol use disorder (Krystal, Cramer, Krol, Kirk, & Rosenheck, 2001; Srisurapanont & Jarusuraisin, 2005) and opioid dependence (Johansson, Berglund, & Lindgren, 2006) and was shown to reduce relapse risk and craving specifically in alcohol use disorder. Consistent with the view that naltrexone reduces the salience of drug cues by strengthening prefrontal activation (Courtney, Ghahremani, & Ray, 2016), we speculate that the beneficial effects of naltrexone on alcohol use and craving might be explained by increased top-down control of DLPFC over striatal wanting signals as a consequence of opioid receptor blockade (but see Nestor et al., 2017). Our results may thus improve the understanding of neural mechanisms underlying pharmacological treatments of dysfunctional wanting in substance use disorders.

Several limitations are worth to be mentioned. First, we did not assess wanting and linking prior to drug administration, such that we cannot control for potential baseline differences in wanting and liking between drug groups. Thus, it remains possible that the non-significant effects of amisulpride on wanting and of naltrexone on liking are caused by such pre-existing baseline differences, or that the sample size was not sufficient to detect these effects in a between-subject design. We also note that the doses for amisulpride and naltrexone might not have been pharmacologically equivalent. In fact, while 50 mg naltrexone produces 95% mu-opioid receptor occupancy (Weerts et al., 2008), 400 mg amisulpride leads to a lower dopamine receptor occupancy of 85% (Lako, van den Heuvel, Knegtering, Bruggeman, & Taxis, 2013), which might be a further reason for why naltrexone showed stronger effects on behavior and brain activation than amisulpride.

Taken together, our findings deepen our understanding of the neurochemical mechanisms mediating the impact of wanting of rewards on behavior. Opioid receptors are involved in the modulation of the strength of inhibitory prefrontal input to the striatum encoding the behavioral relevance of the wanting dimension of rewards. These insights into the interactions between neuroanatomical and neurochemical brain mechanisms implementing wanting-driven approach behavior advance our understanding of the mechanisms underlying pharmacological treatments of substance use disorders.

## Materials and Methods

### Participants

A total of 121 healthy volunteers (58 females; M_age_ = 21.8 years, range = 18-30), recruited from the internal pool of the Laboratory for Social and Neural Systems Research, participated in the study. According to power analysis assuming the effect size from a previous study in our lab (Kahnt, Weber, Haker, Robbins, & Tobler, 2015), 38 participants per group allow detecting a significant effect (alpha = 5%) with a power of 80%. Three participants were excluded from the analysis due to response omissions in more than 30% of all trials in the rating task (see below), two further participants were excluded due to showing excessive head movement (>5mm in one of the six head motion parameters) in the scanner, resulting in a final sample of 116 participants (placebo: N = 39; naltrexone: N = 37; amisulpride: N = 40). Drug groups were matched with regard to age (*p* = 0.40), sex (*p* = 0.34), years of education (*p* = 0.45), and BMI (*p* = 0.29). Participants were screened prior to participation for exclusion criteria including history of brain disease or injury, surgery to the head or heart and neurological or psychiatric diseases (including alcoholism, depression, schizophrenia, bipolar disorders, claustrophobia or Parkinson symptoms). Further exclusion criteria were a severe medical disease such as diabetes, cancer, insufficiency of liver or kidneys, acute hepatitis, high or low blood pressure, any cardiovascular incidences, epilepsy, pregnancy or breastfeeding, past use of opiates or other drugs that may interact with amisulpride or naltrexone (such as stimulants). A drug urine test was performed to rule out illicit drug use prior the test session (amphetamines, barbiturates, buprenorphine, benzodiazepines, cannabis, cocaine, MDMA, methadone and morphine/opiates). All participants provided written informed consent. For their participation, they received 40 Swiss francs per hour. The study was approved by the ethics committee of the canton of Zurich (KEK-ZH-NR2012-0347) and preregistered on www.clinicaltrials.gov (NCT02557984).

### Procedure

Participants received a pill containing either placebo (N = 40), 400 mg amisulpride (N = 41) or 50 mg naltrexone (N = 40) in a randomized and double-blind manner three hours before performance of the experimental tasks. Amisulpride is a selective dopamine D2/D3 receptor antagonist, whereas naltrexone is an unspecific opioid receptor antagonist that acts primarily on the μ- and κ-opioid receptors, with lesser and more variable effects on δ-opioid receptors (Rosenzweig et al., 2002; Weerts et al., 2008). We asked participants to fast for 6h before arrival at the lab. After task completion, participants answered post-experimental questionnaires, which probed whether they thought they had received a drug or placebo, and measured their mood (one rating was not recorded in the placebo group). We determined amisulpride and naltrexone blood plasma levels immediately before and after the behavioral tasks with high-performance liquid chromatography–mass spectrometry in order to control for the pharmacokinetics of the drugs.

### Task design and procedure

Participants performed a task in which they had to rate how much they wanted or liked forty non-consumable everyday items (Weber et al., 2018). Before performing the rating task in the scanner, we familiarized participants with the items by physically presenting all items to them. The rating task was implemented in Matlab (The MathWorks, Natick, MA, United States) and the Cogent 2000 toolbox. We asked participants to rate each item according to how much they wanted to have it, as well as how much they liked the item at that moment. In each trial, participants first saw a cue indicating the type of rating (wanting or liking) (1s), followed by an image of the item (3s), and finally the rating screen (3.5s). Ratings were provided on a continuous scale using a trackball. Trials were separated by a variable intertrial interval (mean 3s). Each item was rated twice for wanting and twice for liking, resulting in 160 trials split into 4 runs before the game (pre-test) and 4 runs after the game (post-test). Between the pre-test and post-test experimental sessions, participants played a game inside the scanner in which they could win the items. The game consisted of a perceptual task in which participants had to indicate whether the item was presented to the left or the right of the midpoint of the screen. Participants won items that they classified correctly. The difficulty of the game was calibrated such that participants won and lost 50% of the items.

### MRI data acquisition and preprocessing

Whole-brain scanning was performed with a Philips Achieva 3T whole-body MRI scanner equipped with an 8-channel head coil (Philips, Amsterdam, the Netherlands). For each of the 8 scanning runs, 227 T2-weighted whole-brain EPI images were acquired in ascending order. Each volume consisted of 33 transverse axial slices, using field of view 192 mm × 192 mm × 108 mm, slice thickness 2.6 mm, 0.7 mm gap, in-plane resolution 2 mm × 2 mm, matrix 96 × 96, repetition time (TR) 2,000 ms, echo time (TE) 25 ms, flip angle 80°. Additionally, a T1-weighted turbo field echo structural image was acquired for each participant with the same angulation as applied to the functional scans (181 slices, field of view 256 mm × 256 mm × 181 mm, slice thickness 1 mm, no gap, in-plane resolution 1 mm × 1 mm, matrix 256 × 256, TR 8.4 ms, TE 3.89 ms, flip angle 8°).

Preprocessing was performed with SPM 12 (www.fil.ion.ucl.ac.uk/spm). The functional images of each participant were motion corrected, unwarped, slice-timing corrected (temporally corrected to the middle image), and co-registered to the anatomical image. Following segmentation, we spatially normalized the data into standard MNI space. Finally, data were smoothed with a 6 mm FWHM Gaussian kernel and high-pass filtered (filter cutoff = 128 seconds).

### Behavioral data analysis

Behavioral data in the rating task were analyzed with mixed general linear models (MGLMs) using the lme4 package in R. The alpha threshold was set to 5% (two-tailed) and p-values were computed using the Satterthwaite approximation. To replicate our previous findings that winning versus losing items has dissociable effects on wanting and liking ratings, we regressed item-specific ratings in the post-test session on fixed-effect predictors for *Judgement* (wanting versus liking), *Item type* (lost versus won), z-transformed item-specific ratings in the pre-test, and all interaction terms (MGLM-1). All these predictors were also modelled as random slopes in addition to participant-specific random intercepts. We also performed separate analyses for wanting and liking ratings (MGLM-2) where post-test item-specific ratings were predicted by *Item type* and *Pre-test ratings*.

To assess drug effects on wanting and liking ratings, we regressed session- and item-specific ratings on fixed-effect predictors for *Drug* (Amisulpride versus placebo and Naltrexone versus placebo), *Judgement, Session* (pre-test versus post-test), and the interaction effects (MGLM-3). All fixed effects varying on the individual level (i.e., *Judgement, Session*, and *Judgement* × *Session*) were also modelled as random effects in addition to participant-specific intercepts. Again, we performed separate analyses for wanting and liking (MGLM-4), which were identical to MGLM-3 but left out all predictors for *Judgement*.

### MRI data analysis

To investigate drug effects on neural activation related to wanting and liking ratings, we computed two GLMs, following previous procedures (Weber et al., 2018). GLM-1 included an onset regressor for the presentation of the current item and the rating bar (duration = 7 s). This onset regressor was modulated by three mean-centered parametric modulators, i.e., mean session-specific and item-specific wanting and liking ratings as well as decision times (to control for choice difficulty). The parametric modulators were not orthogonalized, such that the results for the regressors indicate the unique variance explained by wanting or liking ratings. A separate regressor modelled all items for which no session- and item-specific value could be computed due to response omissions. GLM-2 was identical to GLM-1 with the only difference that it included separate onset regressors for wanting and liking trials, which allowed assessing judgement-specific and judgement-unspecific neural correlates of wanting and linking (Weber et al., 2018). In both models, the regressors were convolved with the canonical hemodynamic response function in SPM. We also added 6 movement (3 translation and 3 rotation) parameters as covariates of no interest.

For statistical analysis, we first computed the following participant-specific contrasts: For GLM-1, we computed parametric contrasts for wanting ratings and liking ratings (independently of judgment type) in GLM-1. For the second-level analysis, we entered the contrast images from all participants in a between-participant, random effects analysis to obtain statistical parametric maps. To investigate the neural correlates of wanting and liking independently of administered drug, we conducted whole-brain second-level analyses using one-sample *t-*tests. To assess drug effects, we employed second-level independent *t*-tests for naltrexone versus placebo as well as amisulpride versus placebo. For these analyses, we report results that survive whole-brain family-wise error corrections at the peak level. For the figures, we set the individual voxel threshold to *p* < 0.001 with a minimal cluster extent of k ≥ 20 voxels. Results are reported using the MNI coordinate system.

### Psychophysiological interaction (PPI) analysis

To examine how our pharmacological manipulations modulated the frontostriatal gating of wanting and liking, we conducted a whole-brain PPI analysis with the striatum as seed region. We defined the seed region by building a sphere (diameter = 6 mm) around the coordinates of wanting-related striatum activation in GLM-2 (MNI coordinates: x = −6, y = 11, z = −1). To create the regressors for the PPI analysis, we first extracted the average time course from the seed region for each individual participant (physiological regressor). We then multiplied the physiological regressor with psychological regressors for (i) wanting ratings on wanting trials, (ii) liking ratings on wanting trials, (iii) liking ratings on liking trials, and (iv) wanting ratings on liking trials. Next, we computed a GLM (PPI-1) that included the interaction terms, the physiological regressor, and the psychological regressors. We also added separate onset regressors for wanting and liking trials as well as movement parameters as regressors of no interest. For the statistical analysis, we computed contrasts for wanting ratings on wanting trials, wanting > liking ratings on wanting trials, liking ratings on liking trials, and finally liking > wanting ratings on liking trials. We submitted these contrasts to a second-level analysis to yield statistical parametric maps with a one-sample t test.

## Acknowledgements

We thank Karl Treiber for expert support with data collection as well as Beatrice Beck Schimmer for medical support.

## Data availability statement

The behavioral data that support the findings of this study are available on Open Science Framework (https://osf.io/6cevt/; link for peer review: https://osf.io/6cevt/?view_only=bf3df44c90604b2e8deec830b0a9fed8).

